# Practical media formulations for rapid growth of *Lactobacillus iners* and other vaginal bacteria

**DOI:** 10.1101/2024.09.13.612917

**Authors:** Daniella Serrador, Jhenielle R. Campbell, Dorothy Cheung, Gelila Shefraw, Rupert Kaul, William W. Navarre

## Abstract

Vaginal microbiome composition is closely tied to host health. A microbiome dominated by specific anaerobes (e.g., *Gardnerella vaginalis*) is termed bacterial vaginosis (BV) and is associated with negative health outcomes, while colonization by *Lactobacillus* species is thought to protect against BV. However, the role of the species *Lactobacillus iners* in vaginal health is controversial, with evidence that some strains may not protect against BV while others do. To better characterize *L. iners* strains, their interactions with vaginal bacteria and human cells need to be investigated *in vitro*, but this has been impeded by the lack of liquid media for rapid *L. iners* growth. We have developed three liquid media formulations for *L. iners* growth: Serrador’s Lactobacilli-adapted Iscove’s Medium (SLIM) which leads to robust *L. iners* growth, a vaginally adapted version of SLIM (SLIM-V) and a chemically defined medium (SLIM-CD). SLIM and SLIM-V lead to dramatically improved *L. iners* growth compared to previously published formulations and support growth of other vaginal bacteria, including *L. crispatus, L. jensenii, L. gasseri* and *G. vaginalis*. SLIM-CD leads to slower growth but could prove useful for characterizing *L. iners* nutrient requirements or metabolite production. A modified version of SLIM-V supports growth of human cervical epithelial cells and provides a base for future co-culture work. Here, we present the formulations of SLIM, SLIM-V and SLIM-CD, and compare the growth of bacterial strains and human cells in the media.

## Introduction

The composition of the vaginal microbiome is closely tied to host health. Colonization by *Lactobacillus* bacterial species, like *L. crispatus, L. gasseri* and *L. jensenii*, is considered optimal while colonization by other anaerobes, like *Gardnerella* and *Prevotella* species, can lead to bacterial overgrowth, resulting in bacterial vaginosis (BV)^1^. BV is the most common cause of vaginal complaints among reproductive-age women, affecting an estimated 23-29% globally^2^. In addition to vaginal symptoms, BV is associated with increased risk of preterm delivery, low birth weight, cervical cancer, and acquisition of several sexually transmitted infections, including HIV and herpes^3-7^. After standard antibiotic treatment, BV will recur in over 50% of patients^8^. A better understanding of the dynamics of the vaginal microbiome are needed to improve treatment and prevention efforts.

*Lactobacillus* species are thought to protect against BV by directly or competitively inhibiting growth of BV-associated bacteria (BVAB)^9^. A significant proportion of vaginal microbiomes are dominated by one of four *Lactobacillus* species: either *L. crispatus, L. iners, L. gasseri*, or *L. jensenii*, with the former two being more prevalent^10-13^. *L. iners* is estimated to be the most prevalent vaginal bacteria globally^14^. Unfortunately, its role in vaginal health is controversial, with some labeling *L. iners*-dominated microbiomes as a ‘transitional’ state, between the other ‘healthy’ *Lactobacillus* and BV^15, 16^. *L. iners*-dominated microbiomes transitioned to BV more frequently on average than those dominated by the other *Lactobacillus*, though notably some *L. iners*-dominated women were stably colonized by *L. iners* while others had frequent BV transitions^17^. Unlike the other *Lactobacillus, L. iners* frequently co-colonizes the vagina along with BVAB during BV^11^. Additionally, *L. iners* produces a pore-forming toxin, inerolysin, capable of lysing human epithelial cells, which has elevated expression in patients with BV^18, 19^. However, there is also evidence that some *L. iners* strains may be able to inhibit growth of BVAB, as some encode lanthipeptides, one of which inhibited growth of *Gardnerella vaginalis* when expressed in *Escherichia coli*^20^.

Several have proposed that some strains of *L. iners* may stably colonize the vaginal microbiome and protect against BV, while others are permissive to BV^14-16^. To investigate this, the impact of *L. iners* strains on BVAB, other *Lactobacillus*, and host cells *in vitro* needs to be determined. While these analyses have been performed with the other *Lactobacillus* species, *L. iners* is frequently omitted^21, 22^. One study examining interactions of *Lactobacillus* and BVAB in co-culture included only a single *L. iners* strain, and another study comparing how supernatant from strains of *Lactobacillus* impacted *G. vaginalis* and *Prevotella bivia* growth excluded *L. iners* completely^21, 22^. This may have been due to the lack of liquid media for robust *L. iners* growth.

*L. iners* does not grow in de Man-Rogosa-Sharpe (MRS) broth, the standard cultivation media for *Lactobacillus* species^23, 24^. Recently, Bloom *et al*. published a medium for *L. iners* growth that consisted of MRS with additional cysteine and glutamine (MRS-CQ)^25^. In MRS-CQ, they found peak *L. iners* growth required 36 hours of incubation^25^. This is suboptimal for co-culture growth assays with other *Lactobacillus*, which grow to high density after just 18 hours in MRS. MRS-CQ was not designed to mimic the native vaginal environment and does not support growth of human cells, which would be useful for creating better models of the vaginal environment. Additionally, MRS is not a chemically defined medium^24^. It is composed of beef extract, yeast extract, and peptone, which is a tryptic meat digest, which vary by batch or manufacturer and make analyzing nutrient requirements or produced metabolites difficult^24^.

To facilitate further *in vitro* characterization of *L. iners*, we have developed three inexpensive and easy to make liquid media formulations for improved *L. iners* growth. Serrador’s *Lactobacillus*-adapted Iscove’s Medium (SLIM) leads to rapid and robust growth of several *L. iners* strains, as well as vaginal *Lactobacillus* and *Gardnerella* strains. *L. iners* growth is slightly improved by a vaginally adapted medium (SLIM-V). A modified version of SLIM-V can be used for culturing cervical epithelial cells. Finally, a fully defined medium (SLIM-CD) grows several *L. iners* strains and will prove useful for metabolite characterization or analyzing the impact of specific nutrients on *L. iners* growth.

## Methods

### Media preparation

To create MRS-CQ, MRS broth (NutriSelect® Basic, Sigma-Aldrich, St. Louis, MO, 69966-500G) with 0.1% TWEEN® 80 (see table 1) was autoclaved before addition of L-Cysteine and L-Glutamine (see table 1) at concentrations used by Bloom *et al* (4 mM and 1.1 mM, respectively)^25^. Media was passed through a 0.2 μm filter (Nalgene™, ThermoFisher, Waltham, MO, 595-5420) and stored at 4 °C until use.

**Table 1:**
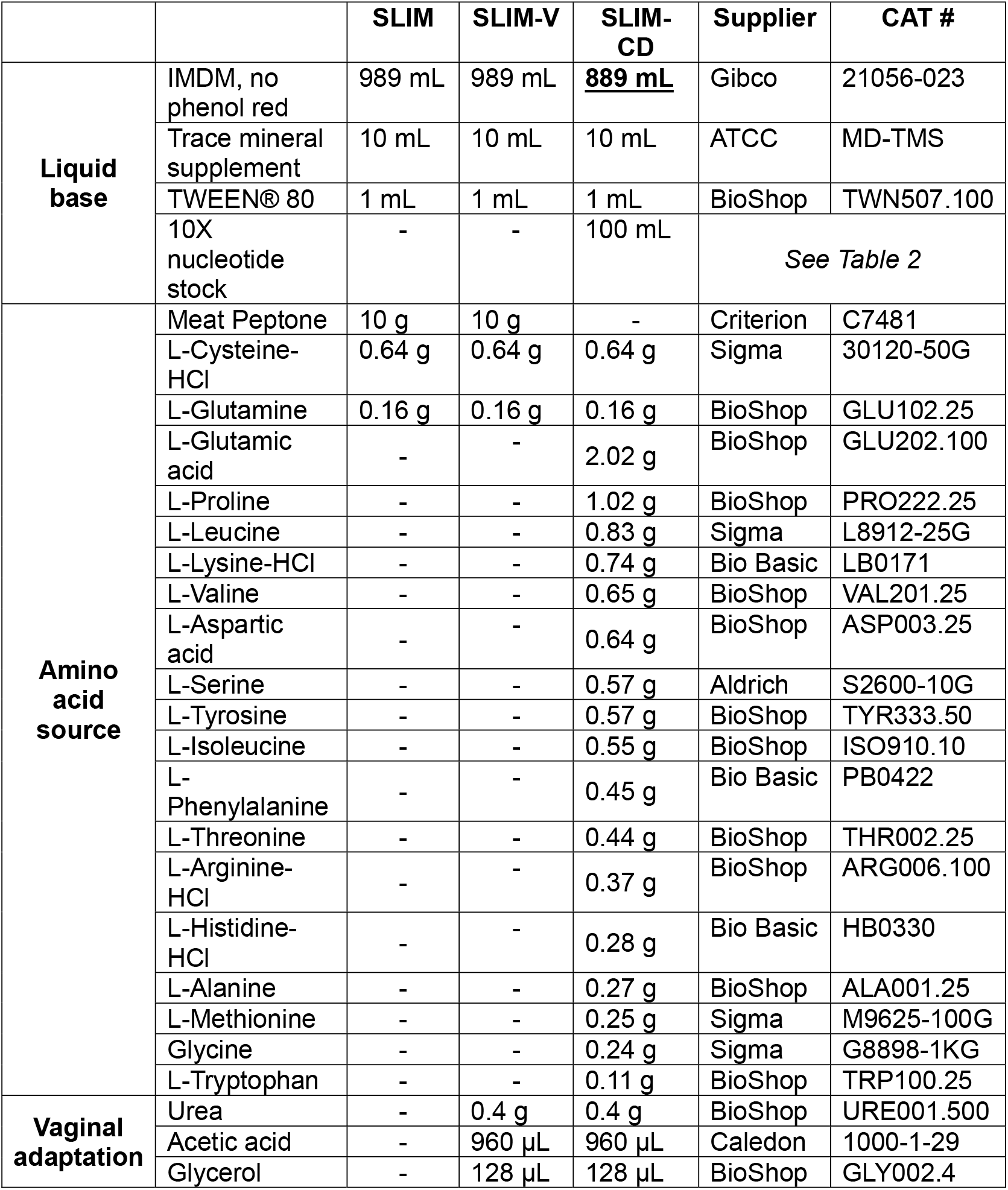
Formulations for SLIM, SLIM-V and SLIM-CD. Recipes to create 1 L of media. (−) indicates component not added. When a component’s amount differs between formulations, values are bolded and underlined.

|  |  | SLIM | SLIM-V | SLIM-CD | Supplier | CAT # |
| --- | --- | --- | --- | --- | --- | --- |
| **Liquid base** | IMDM, no phenol red | 989 mL | 989 mL | <u>**889 mL**</u> | Gibco | 21056-023 |
|  | Trace mineral supplement | 10 mL | 10 mL | 10 mL | ATCC | MD-TMS |
|  | TWEEN® 80 | 1 mL | 1 mL | 1 mL | BioShop | TWN507.100 |
|  | 10X nucleotide stock | - | - | 100 mL | *See Table 2* |  |
| **Amino acid source** | Meat Peptone | 10 g | 10 g | - | Criterion | C7481 |
|  | L-Cysteine-HCl | 0.64 g | 0.64 g | 0.64 g | Sigma | 30120-50G |
|  | L-Glutamine | 0.16 g | 0.16 g | 0.16 g | BioShop | GLU102.25 |
|  | L-Glutamic acid | - | - | 2.02 g | BioShop | GLU202.100 |
|  | L-Proline | - | - | 1.02 g | BioShop | PRO222.25 |
|  | L-Leucine | - | - | 0.83 g | Sigma | L8912-25G |
|  | L-Lysine-HCl | - | - | 0.74 g | Bio Basic | LB0171 |
|  | L-Valine | - | - | 0.65 g | BioShop | VAL201.25 |
|  | L-Aspartic acid | - | - | 0.64 g | BioShop | ASP003.25 |
|  | L-Serine | - | - | 0.57 g | Aldrich | S2600-10G |
|  | L-Tyrosine | - | - | 0.57 g | BioShop | TYR333.50 |
|  | L-Isoleucine | - | - | 0.55 g | BioShop | ISO910.10 |
|  | L-Phenylalanine | - | - | 0.45 g | Bio Basic | PB0422 |
|  | L-Threonine | - | - | 0.44 g | BioShop | THR002.25 |
|  | L-Arginine-HCl | - | - | 0.37 g | BioShop | ARG006.100 |
|  | L-Histidine-HCl | - | - | 0.28 g | Bio Basic | HB0330 |
|  | L-Alanine | - | - | 0.27 g | BioShop | ALA001.25 |
|  | L-Methionine | - | - | 0.25 g | Sigma | M9625-100G |
|  | Glycine | - | - | 0.24 g | Sigma | G8898-1KG |
|  | L-Tryptophan | - | - | 0.11 g | BioShop | TRP100.25 |
| **Vaginal adaptation** | Urea | - | 0.4 g | 0.4 g | BioShop | URE001.500 |
|  | Acetic acid | - | 960 µL | 960 µL | Caledon | 1000-1-29 |
|  | Glycerol | - | 128 µL | 128 µL | BioShop | GLY002.4 |

SLIM is created by adding nutrients to Iscove’s Modified Dulbecco’s Medium (IMDM), a chemically defined, commercially available cell culture medium^26^. To create SLIM, IMDM is supplemented with peptone, L-Cysteine, L-Glutamine, a trace mineral supplement and TWEEN® 80 (Table 1). L-Cysteine and L-Glutamine concentrations were increased as used by Bloom *et al*^25^.

SLIM formulations were prepared by creating the liquid base, then adding an additional amino acid source and, if desired, vaginal adaptations. After preparation, media was passed through a 0.2 μm filter (Nalgene™, ThermoFisher, Waltham, MO, 595-5420) and stored at 4 °C until use.

A vaginal-adapted version of SLIM (SLIM-V) was developed by adding urea, glycerol and acetic acid, which were present in a simulated vaginal fluid developed by Owen & Katz^27^ (Table S1). To create SLIM-V, urea, glycerol, and acetic acid, at concentrations used in the vaginal fluid simulant, were added to SLIM (Table 1).

We observed that *L. iners* grew in SLIM-V when peptone was replaced by a nucleotide supplement (Table 2) and casamino acids (Bio Basic, Markham, ON, CB3060), an acid-hydrolysis of the milk protein casein (Figure S1). To create a chemically defined medium (SLIM-CD), we replaced peptone with the nucleotide supplement and amino acids mimicking the composition of casein supplements reported by Rasmussen *et al*^28^ (Table 1).

**Table 2:**
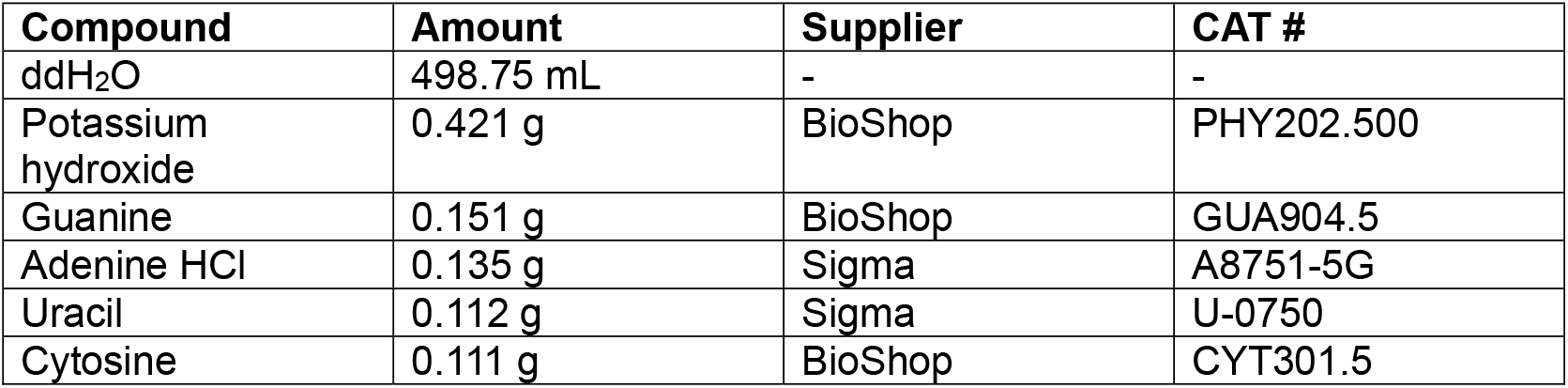
Composition of nucleotide stock. Recipe for 500 mL of 10X stock. Concentrations based on 10X ACGU solution (TekNova, Hollister, CA).

A 10X nucleotide stock was prepared by dissolving nucleotides in ultra-pure water, then passing the solution through a 0.2 μm filter (Nalgene™, ThermoFisher, Waltham, MO, 595-5420). The stock was stored at -20 °C until use. Concentrations were based on 10X ACGU solution (TekNova, Hollister, CA) (Table 2).

### Bacterial cultivation

All bacterial cultivation and growth assays were performed in an anaerobic chamber (AS-580, Anaerobe Systems, Morgan Hill, CA) using a gas mixture of 10% CO_2_, 10% H_2_, balance N_2_ (Linde, Mississauga, ON). Liquid media was allowed to deoxygenate in the chamber for at least 24 hours prior to use, and agar plates were deoxygenated for at least 7 hours prior to use. *L. iners* and *Gardnerella* species were grown on New York City III (NYC III) agar plates^29^. All other lactobacilli, including vaginal *Lactobacillus*, were grown on MRS agar plates. Bacterial stocks were maintained at -80 °C in brain-heart infusion broth (BHI) (Millipore, Darmstadt, Germany, 75917-500G) with 20% glycerol (BioShop, Burlington, ON, GLY002.4). After streaking frozen stocks onto plates, plates were incubated at 37 °C for 48-66 hours.

### Bacterial strains

Experiments included previously reported strains, as well as 6 *L. iners* isolates, 1 *L. crispatus* isolate and 1 *Gardnerella* isolate cultivated from cervicovaginal secretions (Table 3). Cervicovaginal secretion samples were provided by the lab of Dr. Rupert Kaul (Immunology, University of Toronto). In brief, female sex workers in Nairobi, Kenya were recruited through the Sex Worker’s Outreach Program (SWOP) clinics, and they were provided informed, written consent for immune and microbial studies (REB approval in protocol #37046). Consenting participants provided cervicovaginal secretions collected by SoftCup (Evofem, San Diego, CA). Secretions were diluted 10-fold in sterile PBS, then centrifuged at 1730 x g for 10 minutes, after which supernatant was extracted and pellets were resuspended in 500 µL PBS. Both were frozen at -80 °C and transported to the Kaul lab at the University of Toronto for analysis.

**Table 3:** Vaginal bacterial strains used in this study.

| **Species** | **Strain** | **Source** | **Original Study** |
| --- | --- | --- | --- |
| *Lactobacillus iners* | ATCC 55195 | Human vagina | Human Microbiome Project |
| *Lactobacillus iners* | SPIN 2503V10-D | Patient with bacterial vaginosis | Human Microbiome Project |
| *Lactobacillus iners* | NVM020S08 | Cervicovaginal secretions | This study |
| *Lactobacillus iners* | NVM020S14 | Cervicovaginal secretions | This study |
| *Lactobacillus iners* | NVM025S01 | Cervicovaginal secretions | This study |
| *Lactobacillus iners* | NVM025S03 | Cervicovaginal secretions | This study |
| *Lactobacillus iners* | NVM025S06 | Cervicovaginal secretions | This study |
| *Lactobacillus iners* | NVM025S07 | Cervicovaginal secretions | This study |
| *Lactobacillus paragasseri* | JV-V03 | Female urogenital tract | Human Microbiome Project |
| *Lactobacillus gasseri* | EX336960VC01 | Human mid-vaginal wall | Human Microbiome Project |
| *Lactobacillus crispatus* | PSS7772C | Urine of a pregnant woman | Human Microbiome Project |
| *Lactobacillus crispatus* | NVM146S01 | Cervicovaginal secretions | This study |
| *Lactobacillus jensenii* | 269-3 | Human vaginal mucosa | Human Microbiome Project |
| *Lactobacillus jensenii* | EX849587VC03 | Human mid-vaginal wall | Human Microbiome Project |
| *Gardnerella vaginalis* | ATCC 14019 | Vaginal secretions | Human Microbiome Project |
| *Gardnerella* sp. | NVM022C02 | Cervicovaginal secretions | This study |

To isolate bacteria from the cervicovaginal secretions, the pellets were thawed and 10-fold serial dilutions in sterile 1X PBS were created. Dilutions were spread onto agar plates (NYC III for *L. iners* and *Gardnerella* isolations, MRS for *L. crispatus* isolations) and incubated for 48 hours at 37 °C. Single colonies were picked and restruck onto agar plates, then incubated for 48 hours at 37 °C. This was repeated twice, at which point colony morphology for each isolate was consistent, and frozen stocks were created.

Each isolate was identified using PCR with species-specific primers (Table S2). Isolates NVM020S08 and NVM020S14 were isolated from the same vaginal sample, and isolates NVM025S01, NVM025S03, NVM025S06 and NVM025S07 were also isolated from one sample. Thus, it is possible within each set of isolates, there is the same strains cultured multiple times. However, we believe them to be different strains as random amplified polymorphic DNA typing yielded differing banding patterns for these isolates^30^.

### Growth curve measurements

In the anaerobic chamber, for each media type, three 13 mL polypropylene tubes (Sarstedt, Nümbrecht, Germany, 62.515.006) per bacterial strain tested were prepared with 4 mL of the deoxygenated media. Tubes were inoculated with 2-3 bacterial colonies from fresh agar plates, then incubated anaerobically at 37°C. At each timepoint, cultures were removed from the incubator, resuspended, and 200 µL from each culture, as well as media blanks, was aliquoted into a 96-well plate (Sarstedt, Nümbrecht, Germany, 83.3924), after which cultures were returned to the incubator. The 96-well plate was removed from the anaerobic chamber, and optical density at 600 nm (OD_600_) was measured using a SpectraMax Plus 384 Microplate Reader (Molecular Devices, San Jose, CA).

### Cell Culture and Viability Assay

Human endometrial carcinoma cell line (HEC-1-B) was acquired from Dr. Scott Gray-Owen (Molecular Genetics, University of Toronto). The epithelial cells were maintained in DMEM (Wisent, Saint-Jean-Baptiste, QC, 319-005-CL) supplemented with 10% Fetal Bovine Serum (Gibco, Waltham, MO, 12483-020) and 1% Penicillin-streptomycin (Gibco, Waltham, MO, 15140-122). Cells were grown at 37 °C in 5% CO_2._ The viability of the HEC-1-B cells in the developed media were tested using a resazurin assay, where the fluorescence of metabolized resazurin (resorufin) is indicative of the number of viable cells^31^. The HEC-1-B cells were harvested from T25 flasks using a standard trypsinization protocol and passaged in DMEM into a 96-well plate (100µl /well). Following overnight growth (50-60% confluency), the media was removed from the wells and the adherent cells were washed with 1X PBS. Experimental media (SLIM, SLIM-V and SLIM-CD) and control media (DMEM and IMDM) were aliquoted into the appropriate wells in triplicate. A set of wells without cells were reserved as media-only controls. After a 3-hour incubation the cells were imaged using a light microscope (Evos FLoid, ThermoFisher, Waltham, MO, 4471136) before 20 µl of a 0.15 mg/ml resazurin solution was added to each well and incubated for 4 hours. An Infinite M Nano^+^ microplate reader (Tecan, Mannedorf, Switzerland, 30190087) was used to determine cell viability by measuring fluorescence at 560 nm excitation and 590 nm emission.

## Results

### *L. iners* growth in SLIM formulations

For all *L. iners* strains tested, growth was significantly greater in SLIM and SLIM-V compared to MRS-CQ, with all strains but *L. iners* ATCC 55195 showing no growth in MRS-CQ after 66 hours (Figure 1). This is in accordance with previous experiments in our lab, which saw turbid growth in MRS-CQ for *L. iners* SPIN 2503V10-D only after 6 days of incubation (Figure S2). For all strains, growth was slightly (though sometimes not significantly) improved in SLIM-V compared to SLIM (Figure 1). SLIM-CD grew all strains except NVM020S08 and NVM020S14, though peak growth did not occur until 66 hours, unlike SLIM and SLIM-V with peak growth at 24 or 42 hours (Figure 1).

**Figure 1:**
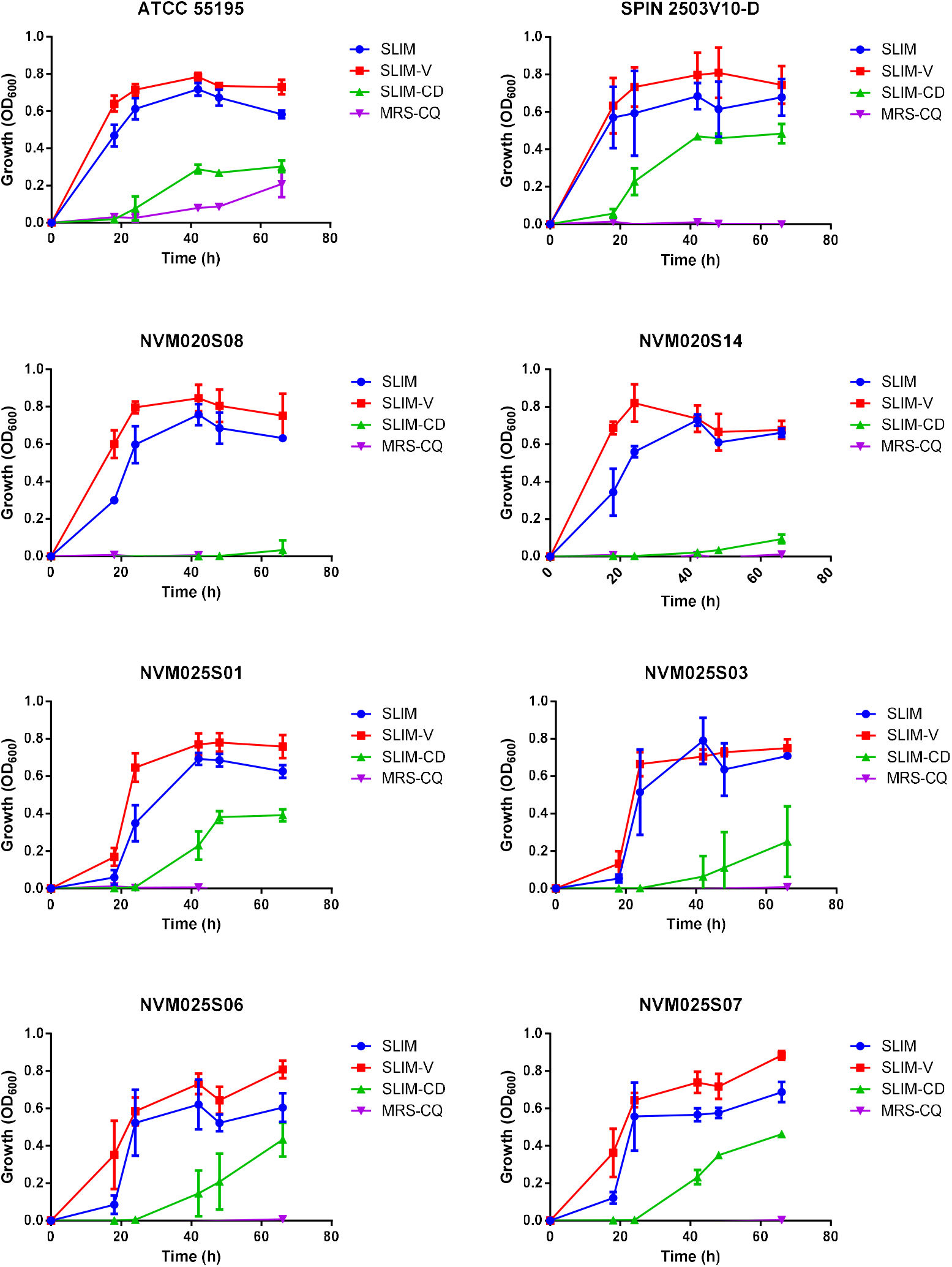
Growth of *L. iners* strains in SLIM, SLIM-V, SLIM-CD and MRS-CQ. OD_600_ measured using SpectraMax Plus 384 Microplate Reader (Molecular Devices, San Jose, CA). n = 3, mean ± SD.

### Growth of other bacteria in SLIM

SLIM and SLIM-V support growth of strains of *L. crispatus* and *L. jensenii*, as well as a strain of *L. gasseri, L. paragasseri, G. vaginalis*, and an unclassified *Gardnerella* species (Figure 2). Growth in SLIM and SLIM-V was similar (Figure 2).

**Figure 2:**
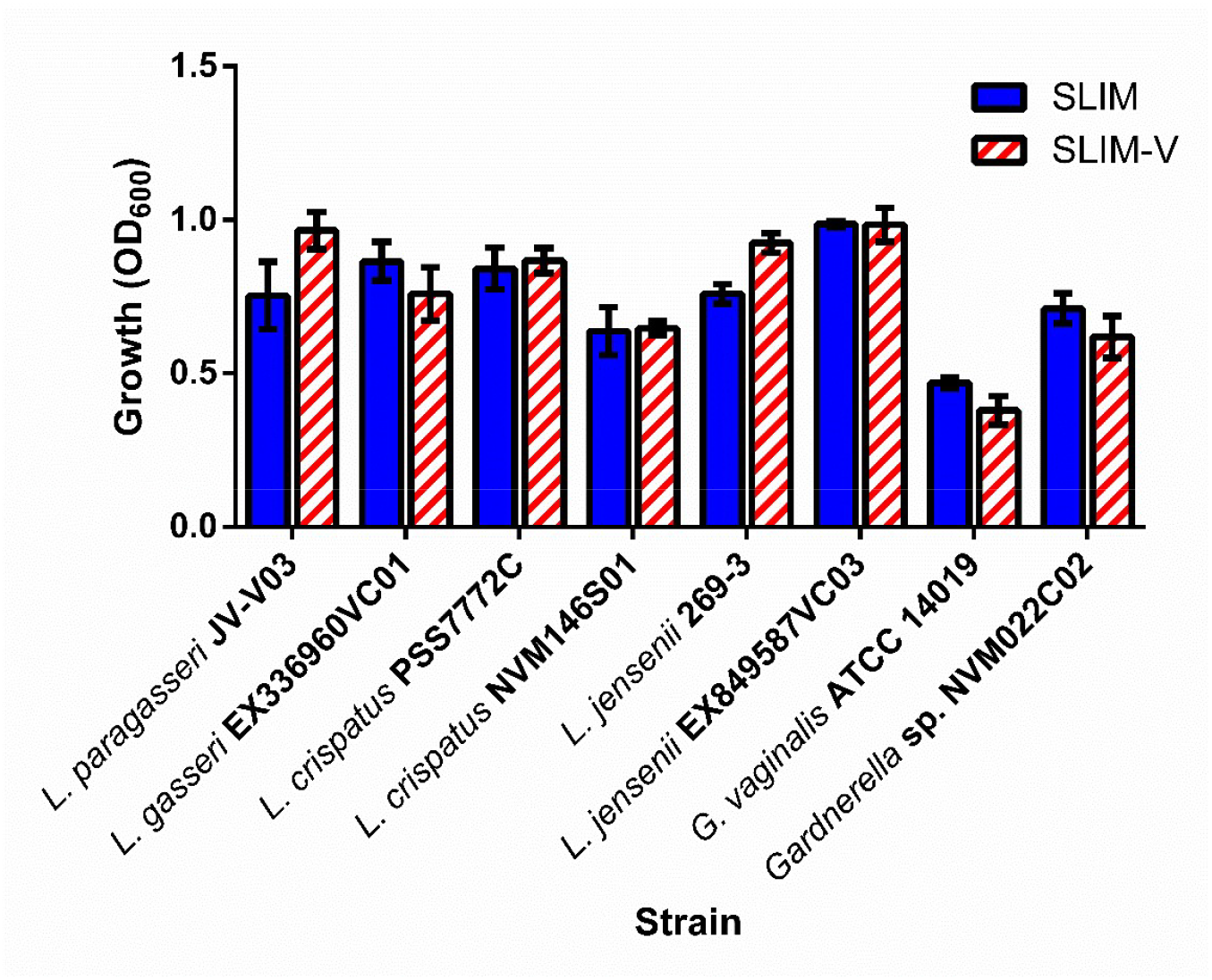
Growth of vaginal bacteria in SLIM and SLIM-V. Growth measured after 24 hours of incubation at 37 °C. OD_600_ measured using SpectraMax Plus 384 Microplate Reader (Molecular Devices, San Jose, CA). n = 3, mean ± SD.

SLIM additionally supports growth of several non-vaginal lactobacilli, including *Limosilactobacillus reuteri, Lactiplantibacillus plantarum, Ligilalactobacillus murinus, Ligilalactobacillus ruminis, Logiolactobacillus coryniformis, Lacticaseibacillus rhamnosus, Companilactobacillus farciminis, Lactobacillus intenstinalis, Lactobacillus psittaci* and *Lactobacillus johnsonii* (Table S3, Figure S3).

### Growth of Human Cells

The HEC-1-B epithelial cells were viable in both SLIM and SLIM-V; however, viability was significantly dampened in SLIM-CD (Figure 3). The inclusion of TWEEN® 80 in all media formulations, except for SLIM-CD, resulted in increased cell viability based on higher levels of resazurin metabolism (Figure 3). However, the morphology of the cells in media formulations containing TWEEN® 80 displayed considerably more vacuoles compared to media without TWEEN® 80 (Figure 4), suggesting that HEC-1-B cells are under stress in the presence of TWEEN® 80. Comparing both viability and morphology, SLIM-V without TWEEN® 80 is the condition most like standard DMEM (Figures 3 & 4).

**Figure 3:**
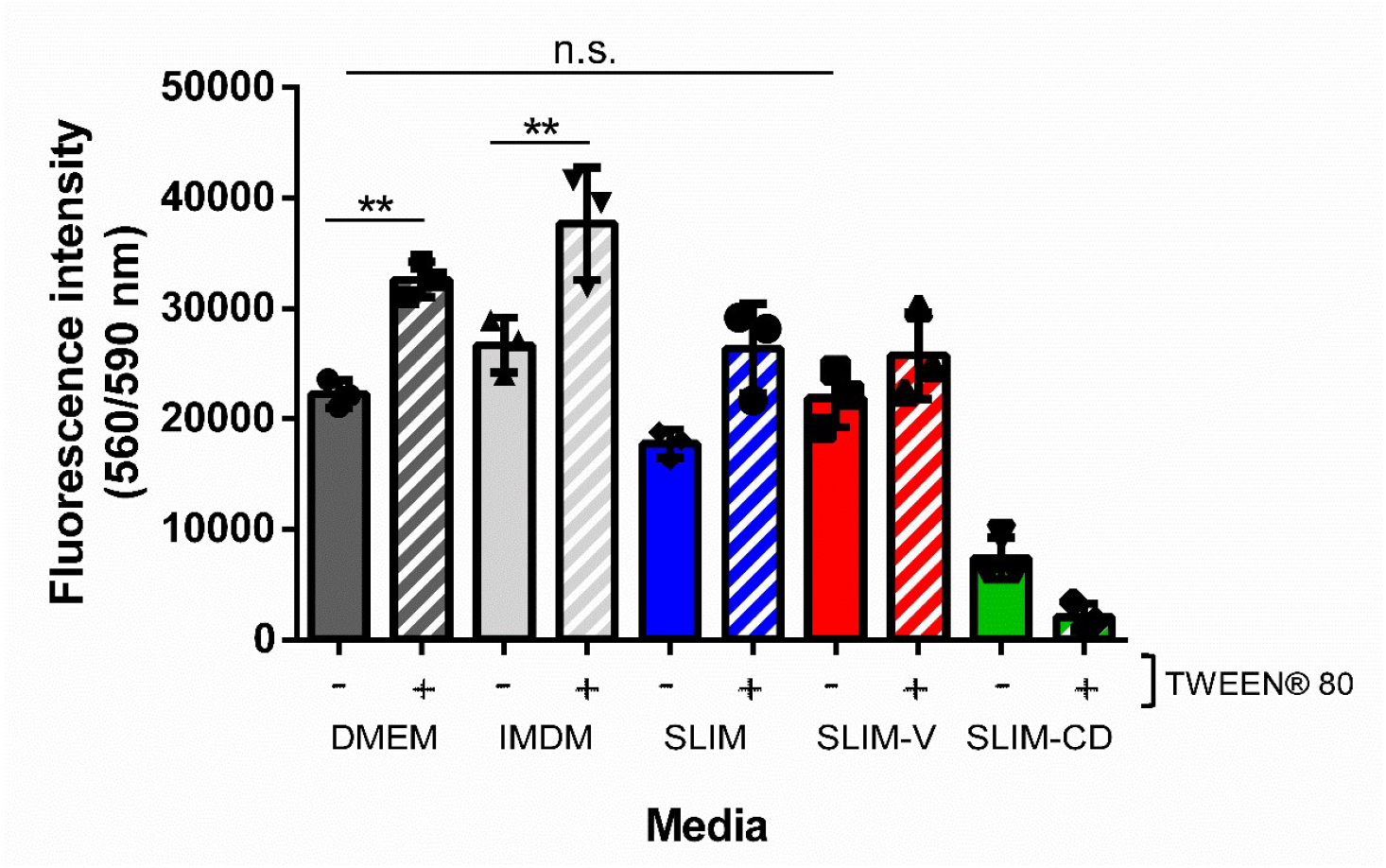
Resazurin-based cell viability of HEC-1-B cells in SLIM formulations. The viability of the cells is directly proportional to the fluorescence intensity of resorufin produced by live cells metabolizing resazurin^31^. Fluorescence intensity of the media-only controls is subtracted from the fluorescence intensity of the appropriate sample wells. Each bar represents the mean ± SD, n=3. Statistical significance was calculated using one-way ANOVA, **\***p < 0.05.

**Figure 4:**
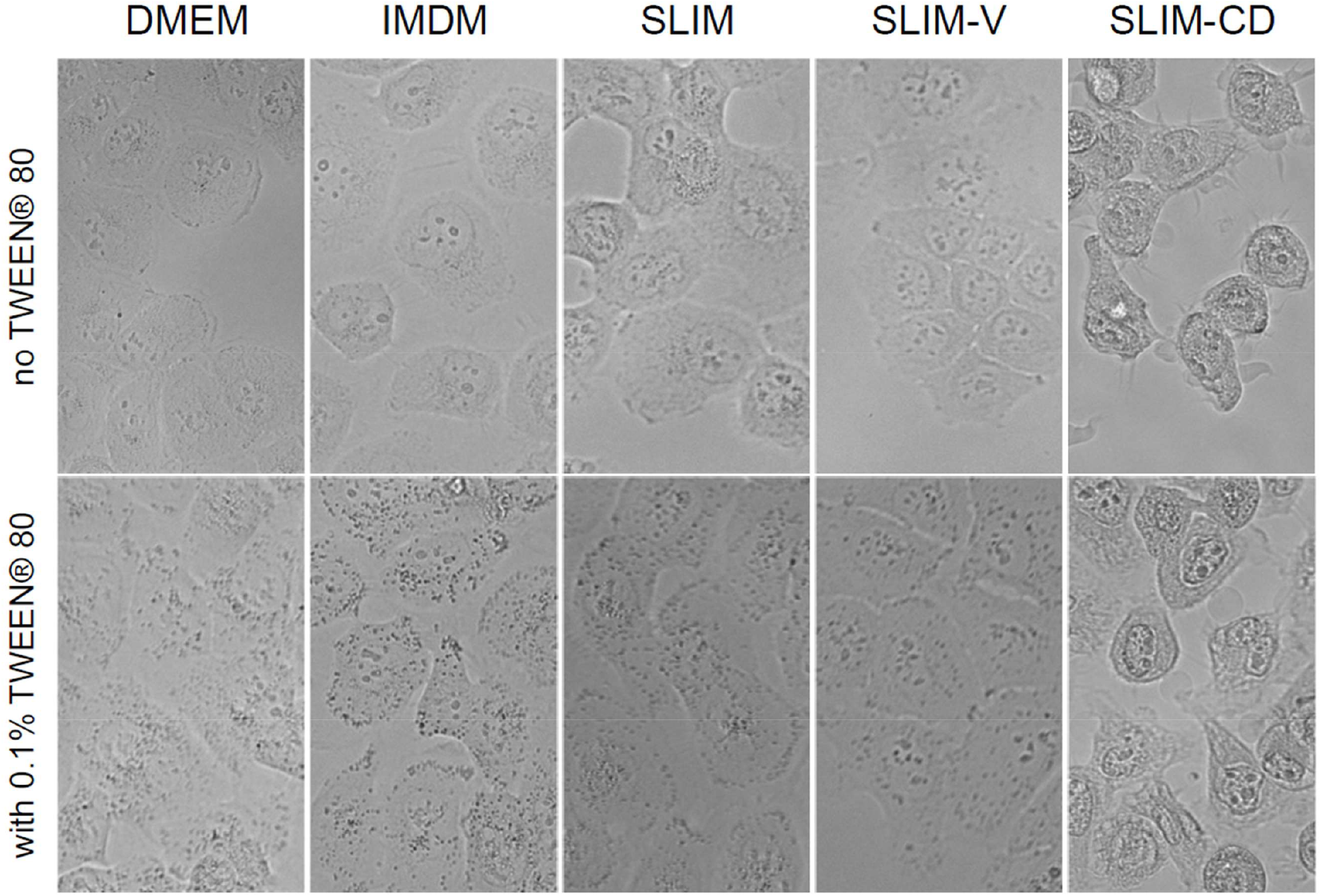
Light microscopy images of HEC-1-B cells in media after 3 hours of incubation.

## Discussion

The role of *L. iners* in the vaginal microbiome is currently poorly understood, in large part due to difficulties in cultivation. Here, we present three media formulations, two leading to robust growth of several *L. iners* strains and other vaginal bacteria, and one that is chemically defined for nutrient and metabolite analysis. These are not minimal media, containing only the nutrients necessary for growth, as we aimed to grow a range of *L. iners* strains as well as other vaginal bacteria and human cells. The trace mineral supplement could likely be replaced with fewer components, and it is probable that not all amino acids added to SLIM-CD are necessary, especially as IMDM already contains a complete set of amino acids. Initial testing suggested that a mixture of several amino acids needed to be added to the IMDM to support growth in SLIM-CD, rather than a single amino acid being required in high amounts.

It is surprising that *L. iners* growth is improved in SLIM compared to MRS-CQ (Figure 1), which is a nutritionally richer medium. There are two potential explanations: that a component of MRS is inhibitory to *L. iners* growth and absent from SLIM, or that components of SLIM that are absent in MRS lead to improved growth. In preliminary testing, *L. iners* ATCC 55195 and SPIN 2503V10-D growth in a 1:1 mixture of MRS-CQ and SLIM was similar to growth in SLIM (Figure S4). This suggests the latter hypothesis is more likely. One possibility is the presence of free amino acids in IMDM, as opposed to amino acids in peptides, in MRS. Bloom *et al*. suggested *L. iners* has a limited capacity to use L-Cysteine from complex sources, which was why additional free L-Cysteine needed to be added to MRS to grow *L. iners* in MRS-CQ^25^. If this is occurring with additional amino acids, IMDM with free amino acids makes a more suitable media base than MRS. Our observed growth in MRS-CQ (Figure 1, Figure S2) was significantly slower than that observed by Bloom *et al*., where growth was seen in 35 hours^25^. This is likely due to differences in cultivation techniques. Larger volumes require more time for dense bacterial growth, and Bloom *et al*. grew 250 µL of culture, while here we grew 5 mL^25^. Additionally, in Bloom *et al*. starting inoculums were bacteria grown on agar plates resuspended in PBS, while here we used 2-3 bacterial colonies^25^.

Growth was slightly improved for both *L. iners* and epithelial cells in SLIM-V compared to SLIM (Figure 1), which was designed to be more similar to the native vaginal environment. Future work could determine the individual impacts of urea, glycerol and acetic acid. One component absent from SLIM-V that is present in the vaginal tract is glycogen, which has been linked to *Lactobacillus* presence and improves *L. crispatus* growth^21, 32^. Future investigations into the impact of glycogen on *L. iners* could use SLIM-V with additional glycogen or create a modified version of SLIM-CD where glycogen is the primary carbon source instead of glucose. While SLIM-CD grew most *L. iners* strains, it did not grow *L. iners* NVM020S08 and *L. iners* NVM020S14 within 66 hours (Figure 1). As both isolates are from the same participant, this could suggest that isolates from the same microbiome share nutrient preferences. It is also possible that both isolates are clonal.

Co-culture experiments with human and bacterial cells are commonly done in a 3-hour window and it is essential to use a media that supports the growth of both organisms and/or does not inhibit their growth. Of the media formulations tested, SLIM-V without TWEEN® 80 is the most comparable to DMEM in supporting uterine HEC-1-B epithelial cells. TWEEN® 80 is reported to damage epithelial cells, so disrupted cell morphology seen in media with TWEEN® 80 (Figure 4) is not unexpected^33^. Cell stress may be increasing metabolic rate, resulting in the elevated resazurin metabolism seen from cells in media with TWEEN® 80 (Figure 3). TWEEN® 80 is necessary for *L. iners* growth, but TWEEN® 80 concentrations could be altered to attempt to reduce epithelial cell stress. Cells are still viable with TWEEN® 80, so SLIM-V could be used for short term assays. SLIM-V provides a base for future co-culture experiments between *L. iners* and vaginal epithelial cells.

## Acknowledgements

We would like to thank the Kaul lab (Immunology, University of Toronto) for the cervicovaginal secretion samples, and especially Rachel Liu for her assistance; Daniela Atere (Molecular Genetics, University of Toronto) for her work characterizing *L. crispatus* NVM146S01; Dr. Lori Frappier’s lab (Molecular Genetics, University of Toronto), specifically Kathy Shire, for assistance with cell culture; and Dr. Leah Cowen’s lab (Molecular Genetics, University of Toronto) for use of the plate readers. This work was funded by an Exploration Grant from the New Frontiers in Research Fund (NFRFE-2021-00522) and the Canada Graduate Scholarships - Master’s program.

## References

1. Mills, B. B. Vaginitis: Beyond the Basics. Obstet. Gynecol. Clin. North Am. 44, 159– 177 (2017).

2. Peebles, K., Velloza, J., Balkus, J. E., McClelland, R. S. & Barnabas, R. V. High Global Burden and Costs of Bacterial Vaginosis: A Systematic Review and MetaAnalysis. Sex. Transm. Dis. 46 (2019).

3. Gravett, M. G., Hummel, D., Eschenbach, D. A. & Holmes, K. K. Preterm Labor Associated With Subclinical Amniotic Fluid Infection and With Bacterial Vaginosis. Obstetrics & Gynecology 67 (1986).

4. Hillier, S. L. et al. Association between bacterial vaginosis and preterm delivery of a low-birth-weight infant. N. Engl. J. Med. 333, 1737–1742 (1995).

5. Gillet, E. et al. Association between bacterial vaginosis and cervical intraepithelial neoplasia: systematic review and meta-analysis. PLoS One 7, e45201 (2012).

6. Gosmann, C. et al. Lactobacillus-Deficient Cervicovaginal Bacterial Communities Are Associated with Increased HIV Acquisition in Young South African Women. Immunity 46, 29–37 (2017).

7. Gallo, M. F. et al. Risk Factors for Incident Herpes Simplex Type 2 Virus Infection Among Women Attending a Sexually Transmitted Disease Clinic. Sex. Transm. Dis. 35 (2008).

8. Bradshaw, C. S. et al. High Recurrence Rates of Bacterial Vaginosis over the Course of 12 Months after Oral Metronidazole Therapy and Factors Associated with Recurrence. J. Infect. Dis. 193, 1478–1486 (2006).

9. France, M., Alizadeh, M., Brown, S., Ma, B. & Ravel, J. Towards a deeper understanding of the vaginal microbiota. Nature Microbiology 7, 367–378 (2022).

10. Lebeer, S. et al. A citizen-science-enabled catalogue of the vaginal microbiome and associated factors. Nature Microbiology (2023).

11. France, M. T. et al. VALENCIA: a nearest centroid classification method for vaginal microbial communities based on composition. Microbiome 8, 166 (2020).

12. Anahtar, M. N. et al. Cervicovaginal Bacteria Are a Major Modulator of Host Inflammatory Responses in the Female Genital Tract. Immunity 42, 965–976 (2015).

13. Fettweis, J. M. et al. Differences in vaginal microbiome in African American women versus women of European ancestry. Microbiology 160, 2272–2282.

14. Holm, J. B., Carter, K. A., Ravel, J. & Brotman, R. M. Lactobacillus iners and genital health: molecular clues to an enigmatic vaginal species. Curr. Infect. Dis. Rep. 25, 67– 75 (2023).

15. Petrova, M. I., Reid, G., Vaneechoutte, M. & Lebeer, S. Lactobacillus iners: Friend or Foe? Trends Microbiol. 25, 182–191 (2017).

16. Vaneechoutte, M. Lactobacillus iners, the unusual suspect. Res. Microbiol. 168, 826–836 (2017).

17. Gajer, G. et al. Temporal Dynamics of the Human Vaginal Microbiota. Sci. Transl. Med. 4, 132ra52 (2012).

18. Ryan, R. et al. Inerolysin, a Cholesterol-Dependent Cytolysin Produced by Lactobacillus iners. J. Bacteriol. 193, 1034–1041 (2011).

19. Macklaim, J. M. et al. Comparative meta-RNA-seq of the vaginal microbiota and differential expression by Lactobacillus iners in health and dysbiosis. Microbiome 1, 12 (2013).

20. Li, L. et al. The First Lanthipeptide from Lactobacillus iners, Inecin L, Exerts High Antimicrobial Activity against Human Vaginal Pathogens. Appl. Environ. Microbiol. 89, e0212322–22. Epub 2023 Feb 27 (2023).

21. Chiara, A. et al. Evaluation of Modulatory Activities of Lactobacillus crispatus Strains in the Context of the Vaginal Microbiota. Microbiology Spectrum 10, 2733 (2022).

22. Happel, A. et al. Exploring potential of vaginal Lactobacillus isolates from South African women for enhancing treatment for bacterial vaginosis. PLOS Pathogens 16, e1008559 (2020).

23. Falsen, E., Pascual, C., Sjödén, B., Ohlén, M. & Collins, M. D. Y. 1. Phenotypic and phylogenetic characterization of a novel Lactobacillus species from human sources: description of Lactobacillus iners sp. nov. International Journal of Systematic and Evolutionary Microbiology, 49, 217–221.

24. De Man, J. C., Rogosa, M. & Sharpe, M. E. A MEDIUM FOR THE CULTIVATION OF LACTOBACILLI. J. Appl. Bacteriol. 23, 130–135 (1960).

25. Bloom, S. M. et al. Cysteine dependence of Lactobacillus iners is a potential therapeutic target for vaginal microbiota modulation. Nature Microbiology 7, 434–450 (2022).

26. Iscove, N. N. & Melchers, F. Complete replacement of serum by albumin, transferrin, and soybean lipid in cultures of lipopolysaccharide-reactive B lymphocytes. J. Exp. Med. 147, 923–933 (1978).

27. Owen, D. H. & Katz, D. F. A vaginal fluid simulant. Contraception 59, 91–95 (1999).

28. Rasmussen, C., Greenwood, M., Kalman, D. & Antonio, J. in 369–407, 2008).

29. Faur, Y. C., Weisburd, M. H., Wilson, M. E. & May, P. S. A new medium for the isolation of pathogenic Neisseria (NYC medium). I. Formulation and comparisons with standard media. Health Lab. Sci. 10, 44–54 (1973).

30. Du Plessis, E. M. & Dicks, L. M. Evaluation of random amplified polymorphic DNA (RAPD)-PCR as a method to differentiate Lactobacillus acidophilus, Lactobacillus crispatus, Lactobacillus amylovorus, Lactobacillus gallinarum, Lactobacillus gasseri, and Lactobacillus johnsonii. Curr. Microbiol. 31, 114–118 (1995).

31. Costa, P., Gomes, A. T. P. C., Braz, M., Pereira, C. & Almeida, A. Application of the Resazurin Cell Viability Assay to Monitor Escherichia coli and Salmonella Typhimurium Inactivation Mediated by Phages. Antibiotics (Basel) 10, 974. doi: 10.3390/antibiotics10080974 (2021).

32. Nunn, K. L. et al. Vaginal Glycogen, Not Estradiol, Is Associated With Vaginal Bacterial Community Composition in Black Adolescent Women. Journal of Adolescent Health 65, 130–138 (2019).

33. Lu, Y. et al. Food Emulsifier Polysorbate 80 Increases Intestinal Absorption of Di-(2-Ethylhexyl) Phthalate in Rats. Toxicol. Sci. 139, 317–327 (2014).

34. You, I. & Kim, E. B. Genome-based species-specific primers for rapid identification of six species of Lactobacillus acidophilus group using multiplex PCR. PLoS One 15, e0230550 (2020).

35. Balashov, S. V., Mordechai, E., Adelson, M. E. & Gygax, S. E. Y. 2. Identification, quantification and subtyping of Gardnerella vaginalis in noncultured clinical vaginal samples by quantitative PCR. Journal of Medical Microbiology, 63, 162–175.

