## Supplemental figures and tables for "Practical media formulations for rapid growth of *Lactobacillus iners* and other vaginal bacteria"

### Supplemental Material

**Table S1: Comparison of simulated vaginal fluid and SLIM components.** Simulated vaginal fluid recipe from Owen & Katz[27]. SLIM concentrations based on IMDM and trace mineral supplement concentrations, as components of peptone are unknown. (–) indicates absent from SLIM.

| **Simulated Vaginal Fluid** | | **Concentration in SLIM (g/L)** | **Reason for Exclusion from SLIM-V** |
|---|---|---|---|
| **Component** | **Concentration (g/L)** | | |
| NaCl | 3.51 | 4.5 | Similar concentration |
| KOH | 1.40 | 0.33 (KCl)<br>0.000076 ($KNO_3$) | Similar concentration |
| $Ca(OH)_2$ | 0.222 | 0.166 ($CaCl_2$) | Similar concentration |
| Glucose | 5.0 | 4.5 | Similar concentration |
| Bovine serum albumin | 0.018 | - | Peptone provides peptide source |
| Lactic acid | 2.00 | - | Produced by *Lactobacillus* |
| Acetic acid | 1.00 | - | **Included** |
| Glycerol | 0.16 | - | **Included** |
| Urea | 0.4 | - | **Included** |

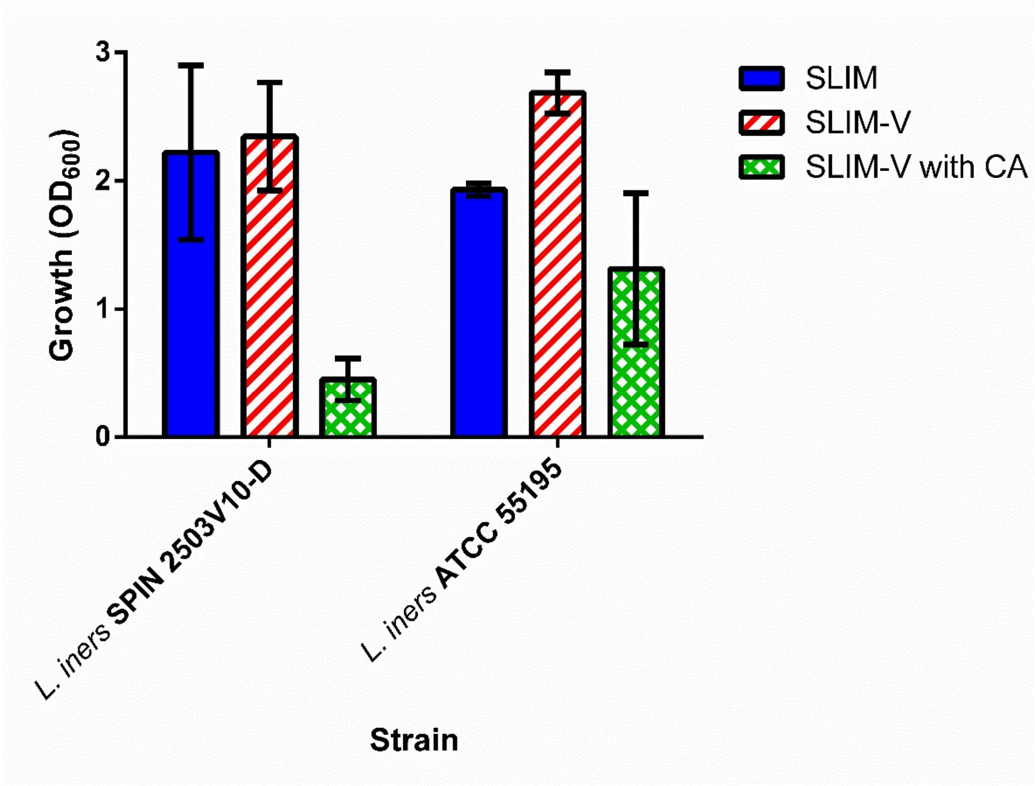


**Figure S1: Growth of *L. iners* and in SLIM, SLIM-V, and SLIM-V with casamino acids.** "SLIM-V with CA" has casamino acids and a nucleotide supplement in lieu of peptone. Growth in 2 mL of media after 24 hours incubation at 37 °C. $OD_{600}$ measured using cuvettes (Sarstedt, Nümbrecht, Germany, 67.742) in Ultrospec 3100 Pro (Amersham Biosciences, Piscataway, NJ). n = 3, mean ± SD.

**Table S2: Primers for identification of *L. crispatus*, *L. iners*, and *Gardnerella* isolates.** Primers target genes unique to each species/genus and conserved among strains of that species/genus.

| Species | Target Gene | Forward | Reverse | Source |
|---|---|---|---|---|
| *L. crispatus* | Hypothetical protein | TGGCGAAGAG ACACCAATATC | TGACGTAACG CATGATGAAT | You & Kim, 2020[34] |
| *L. iners* | Inerolysin | TACTAAGCCTG CACAAGC | TGCATCAAATA CATCACCTGG | This study |
| *Gardnerella* sp. | Elongation factor Tu | TCCCAACCCCA ACTCACGATCTT | NCGCAAACCAAC NATCTCAACTGG | Balashov *et al.,* 2014[35] |

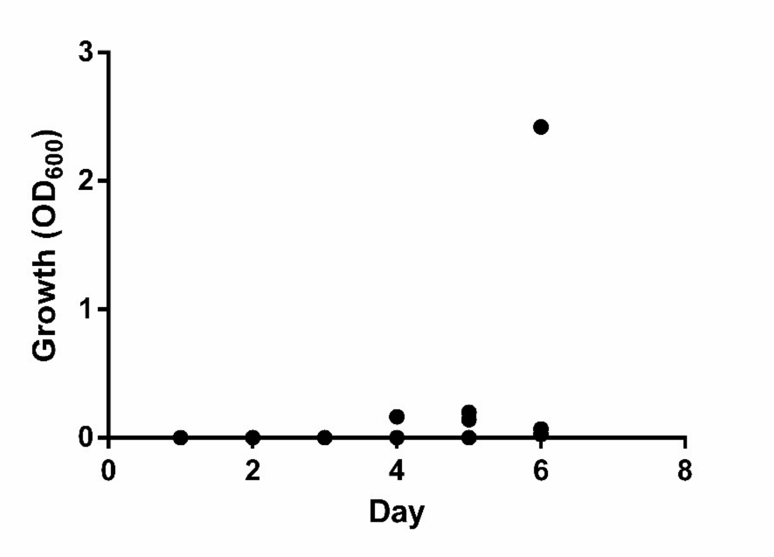


**Figure S2: Growth of *L. iners* SPIN 2503V10-D in MRS-CQ.** 30 tubes with 4 mL of MRS-CQ were inoculated on day 0, and 3 tubes were removed for growth readings each subsequent day until turbid growth was seen. $OD^{600}$ measured using cuvettes (Sarstedt, Nümbrecht, Germany, 67.742) in Ultrospec 3100 Pro (Amersham Biosciences, Piscataway, NJ).

**Table S3: Non-vaginal lactobacilli with growth supported in SLIM.** See Figure S3.

| Species | Strain | Source |
|---|---|---|
| *Limosilactobacillus reuteri* | NM11 | Collection of Inflammation-Associated Mouse Intestinal Bacteria |
| *Limosilactobacillus reuteri* | NM12 | Collection of Inflammation-Associated Mouse Intestinal Bacteria |
| *Lactiplantibacillus plantarum* | DSM 20174 | Pickled cabbage |
| *Ligilalactobacillus murinus* | NM26 | Collection of Inflammation-Associated Mouse Intestinal Bacteria |
| *Ligilalactobacillus murinus* | NM28 | Collection of Inflammation-Associated Mouse Intestinal Bacteria |
| *Ligilalactobacillus ruminis* | DSM 20403 | Bovine rumen |
| *Logiolactobacillus coryniformis* | DSM 20001 | Silage |
| *Lacticaseibacillus rhamnosus* | LMS2-1 | Human gastrointestinal tract |
| *Companilactobacillus farciminis* | DSM 20184 | Sausage |
| *Lactobacillus intenstinalis* | NM61 | Collection of Inflammation-Associated Mouse Intestinal Bacteria |
| *Lactobacillus psittaci* | DSM 15354 | Lung of parrot |
| *Lactobacillus johnsonii* | NM60 | Collection of Inflammation-Associated Mouse Intestinal Bacteria |

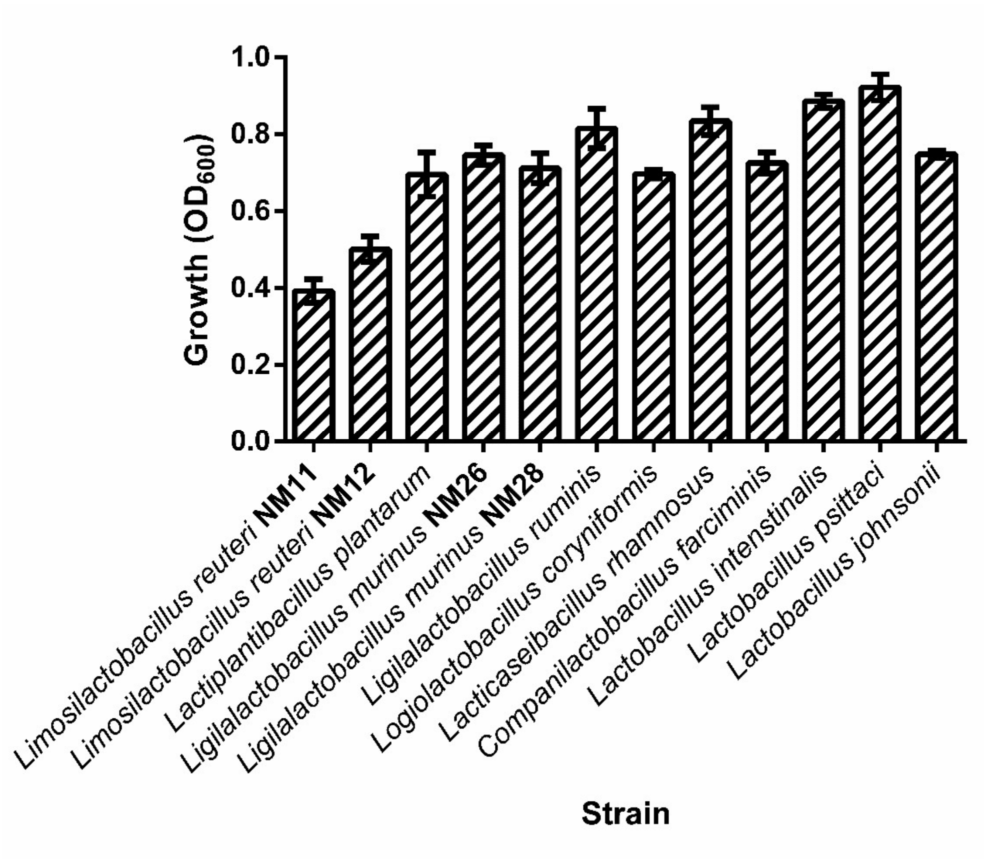


**Figure S3: Growth of other lactobacilli in SLIM.** Growth in 5 mL after 24 hours incubation. $OD_{600}$ measured using SpectraMax Plus 384 Microplate Reader (Molecular Devices, San Jose, CA). n = 3, mean ± SD.

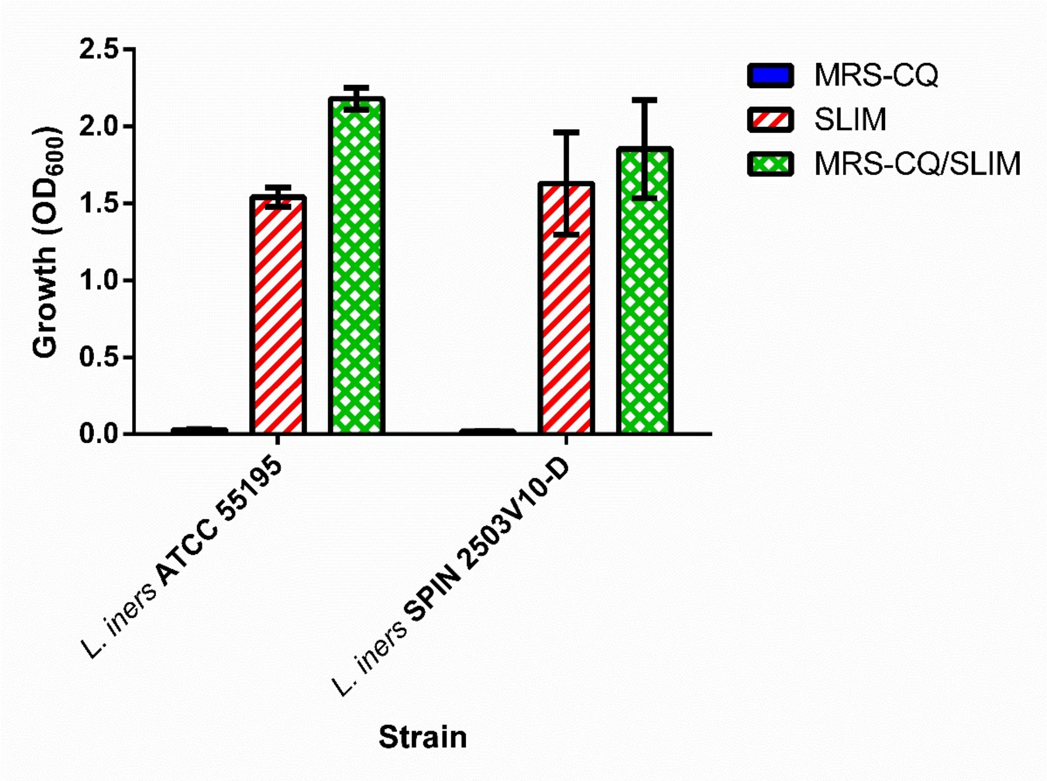


**Figure S4: *L. iners* growth in mixture of MRS-CQ and SLIM.** Growth in MRS-CQ, SLIM, or 1:1 mixture (*MRS-CQ/SLIM*). Growth in 4 mL of media after 48 hours incubation. $OD_{600}$ measured using cuvettes (Sarstedt, Nümbrecht, Germany, 67.742) in Ultrospec 3100 Pro (Amersham Biosciences, Piscataway, NJ). n = 3, mean ± SD.
